# Two uptake hydrogenases differentially interact with the aerobic respiratory chain during mycobacterial growth and persistence

**DOI:** 10.1101/769216

**Authors:** Paul R.F. Cordero, Rhys Grinter, Kiel Hards, Max J. Cryle, Coral G. Warr, Gregory M. Cook, Chris Greening

## Abstract

Aerobic soil bacteria metabolize atmospheric hydrogen (H_2_) to persist when nutrient sources are limited. This process is the primary sink in the global H_2_ cycle and supports the productivity of microbes in oligotrophic environments. To mediate this function, bacteria possess [NiFe]-hydrogenases capable of oxidising H_2_ to subatmospheric concentrations. The soil saprophyte *Mycobacterium smegmatis* has two such [NiFe]-hydrogenases, designated Huc and Hhy, which belong to different phylogenetic subgroups. Huc and Hhy exhibit similar characteristics: both are oxygen-tolerant, oxidise H_2_ to subatmospheric concentrations, and enhance survival during hypoxia and carbon limitation. These shared characteristics pose the question: Why does *M. smegmatis* require two hydrogenases mediating a seemingly similar function? In this work we resolve this question by showing that Huc and Hhy are differentially expressed, localised, and integrated into the respiratory chain. Huc is active in late exponential and early stationary phase, supporting energy conservation during mixotrophic growth and the transition into dormancy. In contrast, Hhy is most active during long-term persistence, providing energy for maintenance processes when carbon sources are depleted. We show that Huc and Hhy are obligately linked to the aerobic respiratory chain via the menaquinone pool and are differentially affected by respiratory uncouplers. Consistent with their distinct expression profiles, Huc and Hhy interact differentially with the terminal oxidases of the respiratory chain. Huc exclusively donates electrons to, and possibly physically associates with, the proton pumping cytochrome *bcc-aa*_3_ supercomplex. In contrast, the more promiscuous Hhy can also provide electrons to the cytochrome *bd* oxidase complex. These data demonstrate that, despite their similar characteristics, Huc and Hhy perform distinct functions during mycobacterial growth and survival.

## Introduction

Earth’s soils consume vast amounts of hydrogen (H_2_) from the atmosphere (1, 2). Over the past decade, research by a number of groups has revealed that this net H_2_ consumption is mediated by aerobic soil bacteria (3–8). Based on this work, it has been established that gas-scavenging bacteria are the major sink in the global H_2_ cycle, responsible for the net consumption of approximately 70 million tonnes of H_2_ each year and 80 percent of total atmospheric H_2_ consumed (6, 9–11). In addition to its biogeochemical importance, it is increasingly realised that atmospheric H_2_ oxidation is important for supporting the productivity and biodiversity of soil ecosystems (12–20). This process is thought to play a key role under oligotrophic conditions, where the majority of microbes exist in a non-replicative, persistent state (14, 21). As the energy requirements for persistence are approximately 1000-fold lower than for growing cells (22), the energy provided by atmospheric H_2_ can theoretically sustain up to 10^8^ cells per gram of soil (23).

The genetic basis of atmospheric H_2_ oxidation has largely been elucidated. Two distinct subgroups of hydrogenase, namely the group 1h and 2a [NiFe]-hydrogenases, are known to oxidise H_2_ to subatmospheric concentrations (3, 5, 24). The operons for these hydrogenases minimally encode the hydrogenase large subunit containing the H_2_-activating catalytic centre, the hydrogenase small subunit containing electron-relaying iron-sulfur clusters, and a putative iron-sulfur protein hypothesised to have a role in electron transfer (23, 25–27). Additional operons encode the maturation and accessory proteins required for hydrogenase function (13, 27). Increasing evidence suggests that hydrogenases capable of atmospheric H_2_ oxidation are widely encoded in soil bacteria. Representatives of three dominant soil phyla, Actinobacteriota, Acidobacteriota, and Chloroflexota, have been experimentally shown to oxidize atmospheric H_2_ (3–5, 8, 24, 28, 29). Moreover, genomic and metagenomic studies indicate that at least 13 other phyla possess hydrogenases from lineages known to support atmospheric H_2_ oxidation (13, 14, 19, 30).

The saprophytic soil actinobacterium *Mycobacterium smegmatis* has served as a key model organism for these studies (24, 27, 31). In *M. smegmatis*, H_2_ oxidation has been shown to be solely mediated by two oxygen-tolerant hydrogenases: the group 2a [NiFe]-hydrogenase Huc (also known as Hyd1 or cyanobacterial-type uptake hydrogenase) and the group 1h [NiFe]-hydrogenase Hhy (also known as Hyd2 or actinobacterial-type uptake hydrogenase) (13, 27). These enzymes belong to distinct phylogenetic subgroups and their large subunits share less than 25% amino acid identity (13). Despite this, Huc and Hhy display striking similarities. Both enzymes oxidise H_2_ to subatmospheric concentrations under ambient conditions (24) and both appear to be membrane-associated despite the lack of predicted transmembrane regions (24). Both Huc and Hhy are reported to be upregulated during stationary phase in response to both carbon and oxygen limitation (27). Consistently, Huc and Hhy deletion mutants show reduced growth yield and impaired long-term survival, suggesting that atmospheric H_2_ oxidation supports energy and redox homeostasis (24, 31, 32). Nevertheless, some evidence suggests that these enzymes are not redundant. For reasons incompletely understood, significant survival phenotypes are observed for both single and double mutants (24, 27). In whole cells, the enzymes also exhibit distinct apparent kinetic parameters, with Hhy having higher affinity but lower activity for H_2_ compared to Huc (24).

It remains to be understood if and how the hydrogenases of *M. smegmatis* are integrated into the respiratory chain. As an obligate aerobe, *M. smegmatis* depends on aerobic heterotrophic respiration to generate proton-motive force and synthesize ATP for growth (33). *M. smegmatis* possesses a branched respiratory chain terminating in one of two terminal oxidases, the cytochrome *bcc-aa*_*3*_ supercomplex or the cytochrome *bd* oxidase (34). The proton pumping cytochrome *bcc-aa*_*3*_ oxidase is the more efficient of these two complexes, leading to the efflux of 6 H^+^ ions per electron pair received, and is the major complex utilised during aerobic growth (34, 35). The non-proton pumping cytochrome *bd* complex is less efficient, resulting in the transport of 2 H^+^ ions per electron pair, but is predicted to have a higher affinity for O_2_ and is important during non-replicative persistence (36, 37). In actively growing *M. smegmatis*, electrons entering the respiratory chain are derived from heterotrophic substrates, and are donated to the respiratory chain by NADH largely via the non-proton pumping type II NADH dehydrogenase NDH-2 and succinate via the succinate dehydrogenase SDH1 (34). While *M. smegmatis* is strictly heterotrophic for replicative growth, it was recently demonstrated that it is able to aerobically respire using carbon monoxide (CO) at atmospheric concentrations during carbon-limited persistence through the actions of a carbon monoxide dehydrogenase (38). It has likewise been predicted that Huc and Hhy support survival during persistence by providing electrons derived from H_2_ to the respiratory chain (24, 27). However, these studies were correlative and it remains to be definitively demonstrated that H_2_ serves as a respiratory electron donor in this organism.

In this work, we addressed these knowledge gaps by comprehensively studying Huc and Hhy during different stages of mycobacterial growth and persistence. We show that Huc and Hhy are differentially expressed throughout growth and persistence and form distinct interactions with the membrane. In addition, we show both Huc and Hhy are obligately linked to the respiratory chain via the menaquinone pool, but form distinct interactions with the terminal oxidases. These data demonstrate that H_2_ oxidation in *M. smegmatis* provides electrons to the respiratory chain for mixotrophic growth via Huc and to energise persistence via Hhy. These findings represent a significant advance in our understanding of the role of high affinity hydrogenases in bacterial metabolism.

## Results and Discussion

### Mycobacterial hydrogenases are differentially expressed and active during growth and persistence

Previous work investigating the activity of Huc and Hhy in *M. smegmatis* showed they are induced in batch culture upon exhaustion of carbon sources (24). However, we lack a high-resolution understanding of the expression and activity of Huc and Hhy during mycobacterial growth and persistence. To address this question, we quantified Huc and Hhy gene expression using qPCR at different growth phases in batch liquid cultures. In exponentially growing, carbon replete cells (OD_600_ 0.3 and OD_600_ 1.0 cultures), transcript levels for *hucL* (the Huc large subunit) were relatively low; however, as carbon sources became exhausted *hucL* expression increased, with maximum expression observed at 1-day post-OD_max_ (OD_600_ ~3.0). Subsequently, at 3-days post-OD_max_ (OD_600_ ~3.0) as cells endured prolonged carbon-limitation, expression levels of *hucL* declined significantly (**Figure 1a**). In contrast, expression of *hhyL* (the Hhy large subunit) remained low during exponential growth, before rapidly increasing by 57-fold when cells reached carbon-limited stationary phase and remained at high levels into late stationary phase (**Figure 1b**).

**Figure 1.**
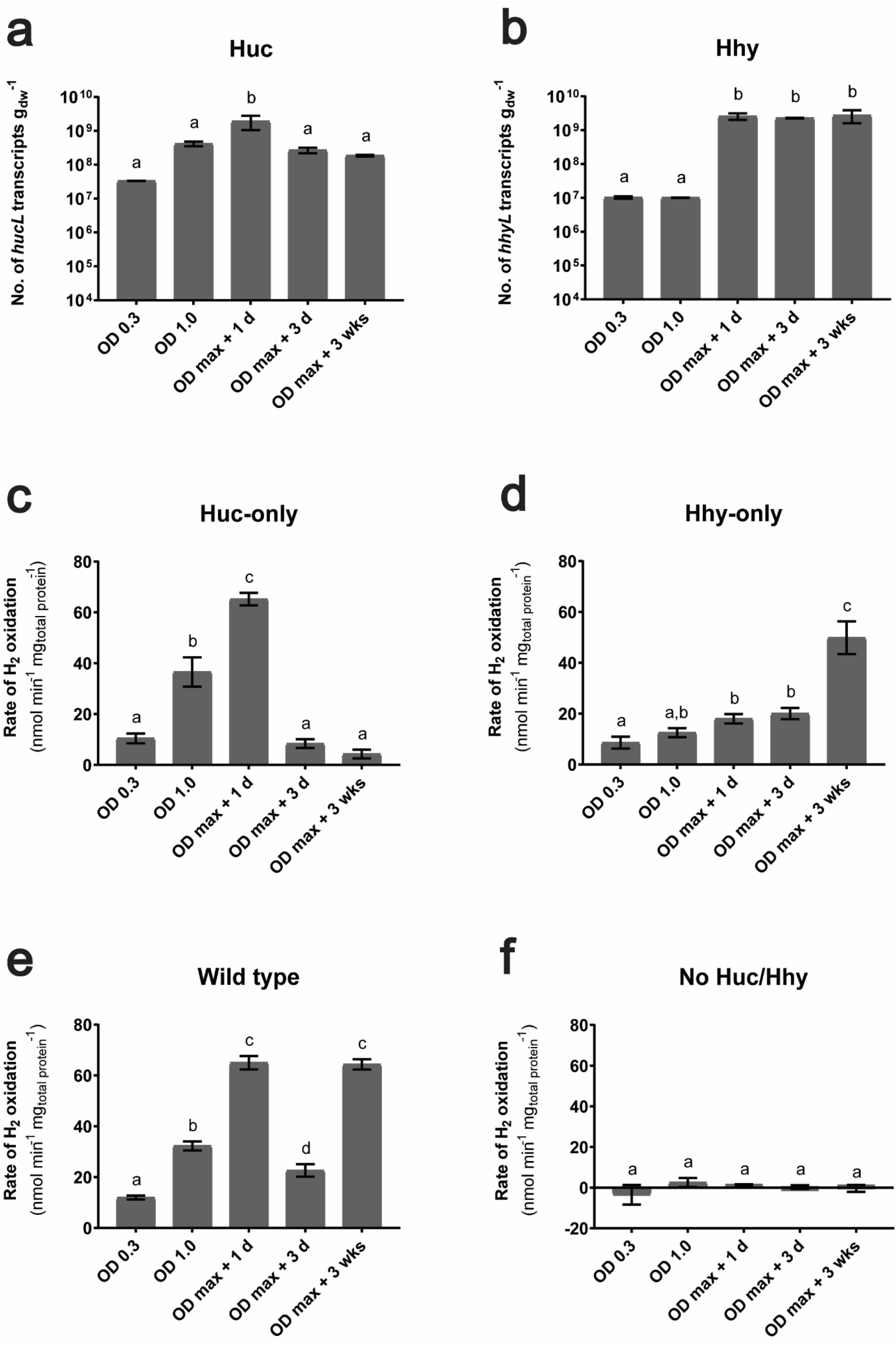
Differential expression and activity of Huc and Hhy. Normalized number of transcripts of the large subunit gene of (**a**) Huc (*hucL*) and (**b**) Hhy (*hhyL*) in wild-type cultures. Cultures were harvested during either carbon-replete conditions, i.e. OD_600_ 0.3 and OD_600_ 1.0, or carbon-limited conditions, i.e. 1 day post-OD_max_ (OD_600_ ~3.0), 3 day post-OD_max_, and 3 weeks post-OD_max_. Absolute transcript levels were determined through qRT-PCR and normalized to the housekeeping gene *sigA*. Rates of H_2_ oxidation of whole cells of (**c**) Huc-only, (**d**) Hhy-only, (**e**) wild-type, and (**f**) no Huc/Hhy (triple hydrogenase deletion) strains of *M*. *smegmatis*. Activities were measured amperometrically using a hydrogen microelectrode under carbon-replete and carbon-limited conditions. All values labelled with different letters are statistically significant based on one-way ANOVA.

Next, we determined the rate of H_2_ oxidation of wild-type *M. smegmatis* and mutant strains containing only Huc or Hhy at different stages of growth and persistence in liquid batch culture. H_2_ oxidation rates in the Huc-only strain correlated well with gene expression levels; levels of H_2_ oxidation were relatively low during early exponential growth (OD_600_ 0.3) and increased during late exponential phase (OD_600_ 1.0), before peaking at 1-day post-OD_max_ and declining rapidly thereafter (**Figure 1c**). The rapid decline in transcript levels and activity of Huc during stationary phase suggests tight regulation of this enzyme. In contrast, the activity of Hhy was low during exponential growth (OD_600_ 0.3 and 1.0), increased slightly at 1- and 3-days post-OD_max_, and increasing markedly during prolonged persistence, with high levels of activity observed at 3-weeks post-_ODmax_ (**Figure 1d**). A notable lag was observed between the increase of transcript levels and Hhy activity during stationary phase, suggesting post-transcriptional regulation of this hydrogenase. The H_2_ oxidation activity profile of the wild-type strain in these assays are the same as the sum of the activity of Huc and Hhy only mutants, confirming that Huc and Hhy are functioning normally in the mutant background (**Figure 1e**). Additionally, a mutant strain lacking both Huc and Hhy did not consume H_2_, confirming that Huc and Hhy are solely responsible for H_2_ oxidation (**Figure 1f**).

These data provide a clear picture of the differential regulation of Huc and Hhy hinted at by previous studies (27). Huc is expressed by *M. smegmatis* during the transition from growth to persistence, allowing cells to grow mixotrophically on atmospheric H_2_ and, where available, higher concentrations produced through abiotic or biotic processes (e.g. fermentation, nitrogen fixation) (39). Subsequently, as cells commit to persistence due to carbon starvation, Hhy is expressed and supplies energy from atmospheric H_2_ to meet maintenance needs.

### Mycobacterial hydrogenases differentially associate with the membrane, with Huc potentially forming a supercomplex with the cytochrome bcc-aa_3_ oxidase

In order to directly attribute the H_2_ oxidation activity in our cellular assays to Huc and Hhy, we separated cell lysates of wild-type and hydrogenase mutant strains using native-PAGE and detected hydrogenase activity by zymographic staining (**Figure 2a**). A high molecular weight species exhibiting H_2_ oxidation activity was detected at 1-day post-OD_max_ in wild-type and Huc-only cultures, but not in the Hhy-only strain. We determined the size of this high-MW species to be >700 kDa *via* blue native-PAGE (**Figure S1**). In contrast, at 3-days post-OD_max_, a low molecular weight H_2_-oxidising species was present in wild-type and Hhy-only cultures, but was absent from the Huc-only strain (**Figure 2a**). These high and low molecular weight bands from the wild-type strain were excised and proteins present were identified by mass spectrometry. The high-MW band yielded peptides corresponding to Huc, while the low-MW band yielded peptides corresponding to Hhy (**Table S1**). These data correlate well with the activity of Huc and Hhy observed in our cellular assays, confirming Huc is the dominant hydrogenase during the transition from growth to dormancy and Hhy is more active during prolonged persistence.

**Figure 2.**
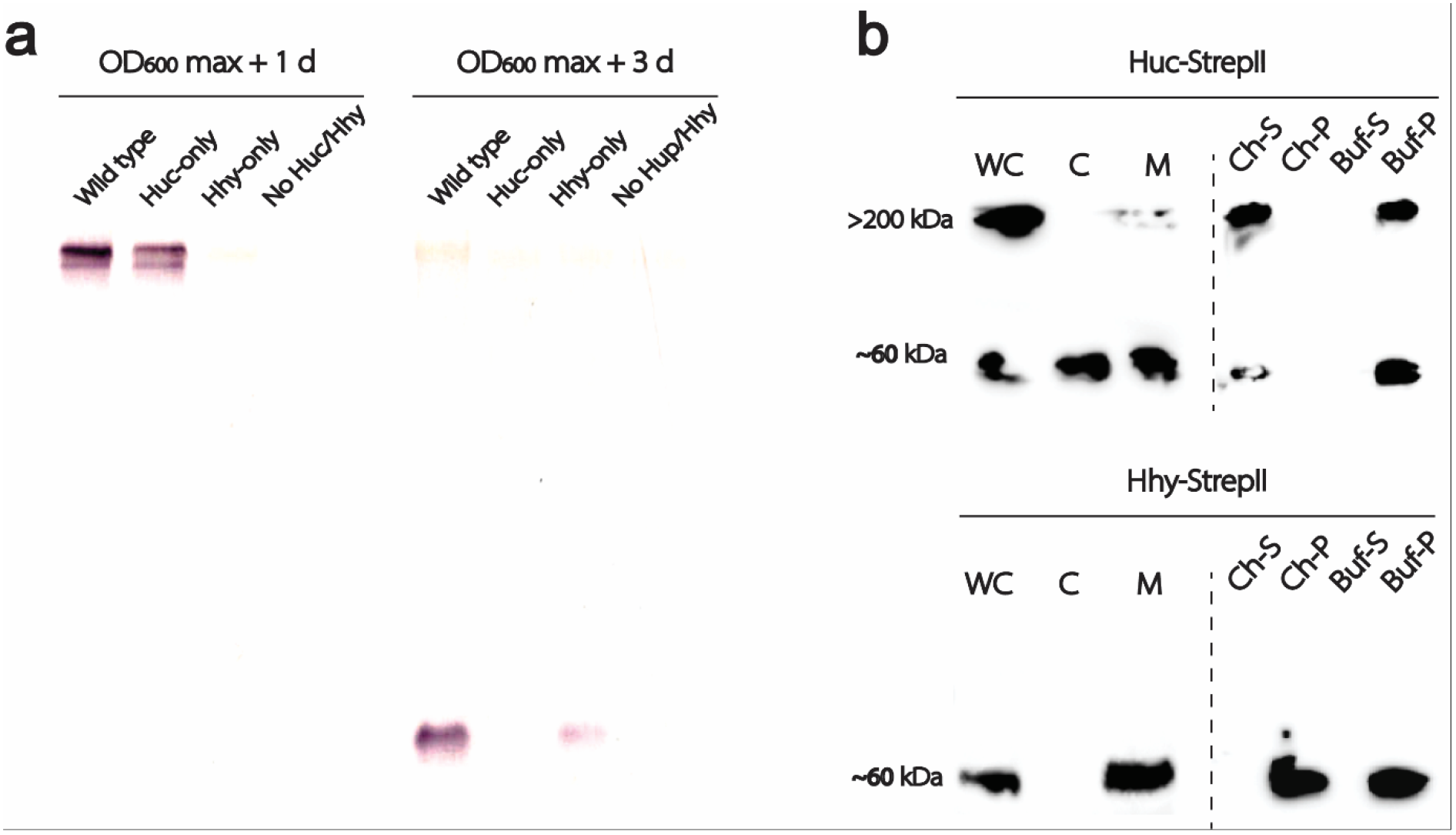
Activity and physical association of Huc and Hhy in cell extracts. (**a**) Differential native activity staining of Huc and Hhy in whole-cell lysates of different *M. smegmatis* strains harvested at 1-day post-OD_max_ (OD_600_ ~3.0) and 3-day post-OD_max_ (OD_600_ ~3.0). (**b**) Localisation of StrepII-tagged Huc (Huc-StrepII) and Hhy (Hhy-StrepII) in different cellular fractions by western blot (left of dotted lines): WC – whole-cell lysates; C – cytosol; M – membrane. Huc-StrepII was harvested at 1-days post-OD_max_ (OD_600_ ~3.0) and Hhy-StrepII 3-days post-OD_max_ (OD_600_ ~3.0). Membranes containing Huc-StrepII and Hhy-StrepII solubilised in 5% sodium cholate at 22 °C for 3 h (right of dotted line). Huc-StrepII and Hhy-StrepII in the cholate-soluble (Ch-S) and cholate-insoluble fractions (Ch-P) are visualized on western blots. Solubilisation controls incubated under identical conditions minus cholate are shown, supernatant (Buf-S) and pellet (Buf-P).

The difference in size between Huc and Hhy activity observed on the native gel is striking. The slow migration of Huc may be due to the formation of an oligomer containing multiple Huc subunits or other unidentified proteins. To test this hypothesis, we interrogated the mass spectrometry data for likely Huc interacting partners. Intriguingly, components of the cytochrome *bcc-aa*_*3*_ oxidase supercomplex were detected in the Huc sample, with a high probability and coverage, demonstrating they are prevalent in this region of the gel (**Table S1**). It was shown previously that H_2_ oxidation in *M. smegmatis* is oxygen dependent, suggesting that Huc and Hhy activity are obligately linked to respiratory chain (24). As cytochrome *bcc-aa*_*3*_ is a large supercomplex (40), association with Huc could account for the high-MW of Huc activity on the native gel, while placing the hydrogenase in an ideal position to donate electrons to this complex.

Previous work indicated Huc and Hhy were membrane-associated despite lacking obvious transmembrane regions or signal peptides (24). To interrogate the nature of this membrane association, we fractionated cells into lysates, cytosols, and membranes and detected the hydrogenases by western blotting chromosomally StrepII-tagged variants of Huc and Hhy. Two bands corresponding to Huc were detected by western blot in *M. smegmatis* whole cells, with sizes of ~60 and >200 kDa. Upon cell fractionation, the ~60 kDa band was observed in both cytoplasmic and membrane fractions, while the >200 kDa band was only observed in the membrane fraction (**Figure 2b**). Interaction of Huc with the membrane was disrupted by 5% sodium cholate, with both bands partitioning to the soluble phase (**Figure 2b**). In contrast, a single ~60 kDa band corresponding to Hhy was observed in the whole cell lysate and membrane fractions, but was absent from the cytoplasmic fraction. The interaction between Hhy and the cell membrane was not disrupted by the addition of 5% sodium cholate, suggesting it forms a strong interaction with the membrane relative to Huc and implies different mechanisms are responsible for their membrane association **(Figure 2b**). Both Huc and Hhy are predicted to form hetero-tetramers, consisting of two large and small subunits with a molecular weight of >200 kDa (26, 27). This is consistent with the bands observed via western blot, with the >200 kDa species observed for Huc representing an intact tetramer, with the ~60 kDa species representing partial disassociation of this complex.

### Mycobacterial hydrogenases are coupled to the respiratory chain and interact differentially with the terminal cytochrome oxidases

While it is known that O_2_ is required for H_2_ oxidation by Huc and Hhy in *M. smegmatis* (6, 24), it had not been determined whether these enzymes support the reduction of O_2_ through coupling to the respiratory chain. To resolve this question, we amperometrically monitored the H_2_ and O_2_ consumption in carbon-limited *M. smegmatis* cells (3-days post-ODmax) following sequential spiking with H_2_ and O_2_ saturated buffer (Figure 3a). Upon spiking cells with H_2_, oxidation (0.31 μM min^−1^) was observed due to ambient levels of O_2_ present in solution. When O_2_-saturated buffer was subsequently added, the rate of H_2_ oxidation increased markedly (1.2 μM min^−1^) (Figure 3a). In cells initially spiked with O_2_, minimal consumption of O_2_ was observed (0.01 μM min^−1^). However, with the subsequent addition of H_2_, the cells consumed O_2_ at approximately half the rate observed for H_2_ oxidation (0.51 μM min^−1^) (Figure 3a). This rate is consistent with the expected stoichiometry of H_2_-dependent aerobic respiration (H_2_ + ½ O_2_ → H_2_O). These data directly link H_2_ oxidation to O_2_ consumption, providing strong experimental evidence that electrons derived from H_2_ support respiratory reduction of O_2_ in *M. smegmatis*.

**Figure 3.**
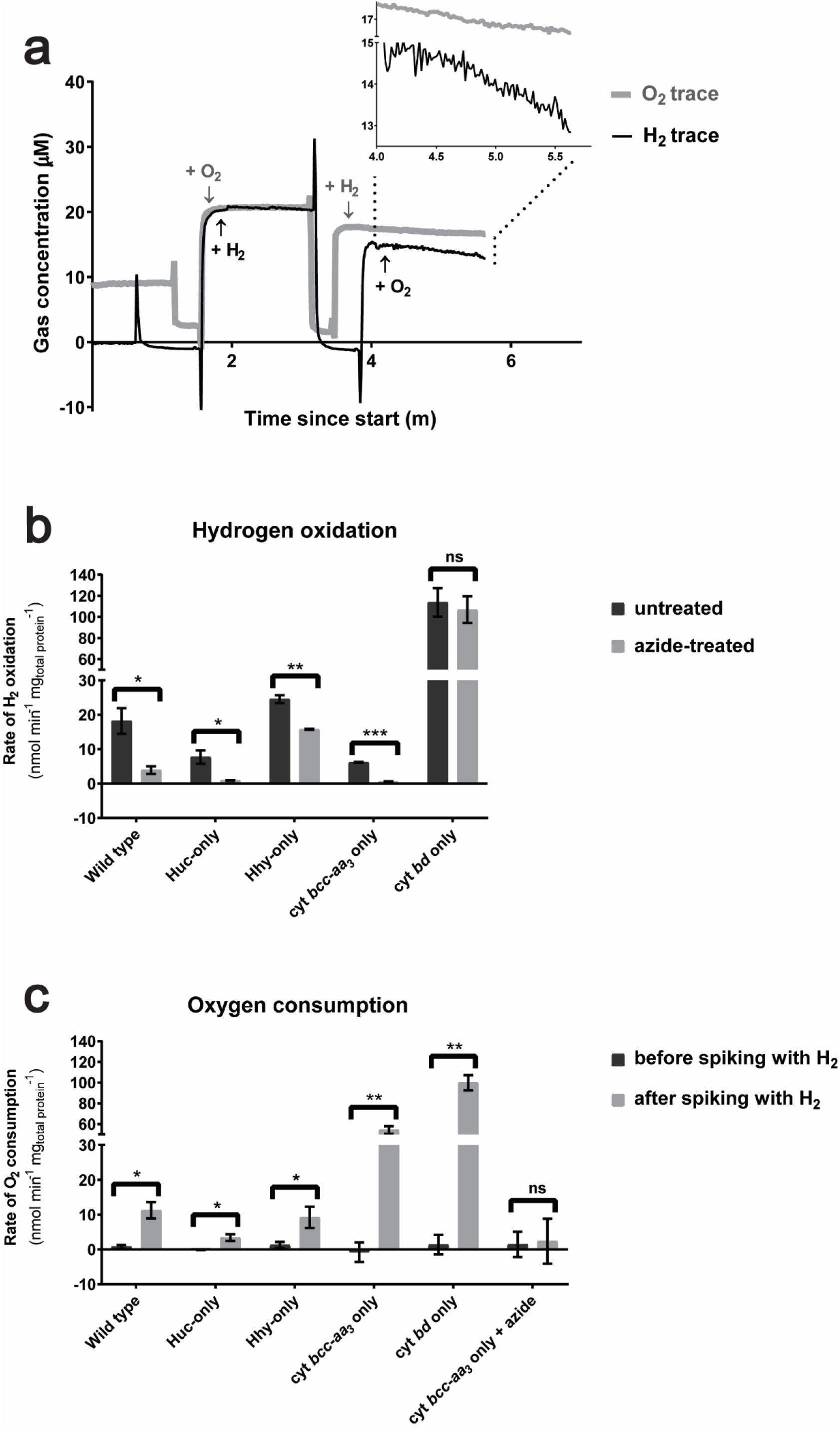
Interaction of Huc and Hhy with the terminal cytochrome oxidases. H_2_ and O_2_ consumption of whole cells from carbon-limited cultures (3 days post OD_max_ ~3.0) of wild-type, hydrogenase, and cytochrome oxidase mutant strains. (**a**) Representative raw electrode traces of H_2_ and O_2_ consumption by carbon-limited wild-type *M. smegmatis* cultures. H_2_ oxidation is dependent in the presence of O_2_ and likewise, O_2_ is not consumed without addition of H_2_ as an electron source. (**b**) The rate of H_2_ uptake by whole cells before and after treatment with the cytochrome oxidase inhibitor zinc azide (250 μM). (**c**) O_2_ consumption in the same set of strains were measured using an oxygen microelectrode, before and after addition of H_2_. Values with asterisks indicate activity rates that are significantly different from the untreated whole cells based on student’s t-test (* p ≤ 0.05; ** p ≤ 0.01; *** p ≤ 0.001; ns – not significant).

Having established that H_2_ oxidation directly supports O_2_ reduction in *M. smegmatis*, we next sought to determine which of the two terminal oxidases were utilized for this process. To achieve this, we monitored the rate of H_2_ oxidation and O_2_ consumption in wild-type, Huc-only, and Hhy-only strains, as well as mutant strains possessing either cytochrome *bcc-aa*_*3*_ or *bd* oxidase as the sole terminal respiratory complex. Given the loss of both terminal oxidases is lethal in *M. smegmatis* (41), we utilised zinc azide, a selective inhibitor of cytochrome *bcc-aa*_*3*_ oxidase (42), to assess the effects of loss of both terminal oxidases on H_2_ oxidation. As expected from our initial experiments at 3-days post-OD_max_ (**Figure 1**), the H_2_ oxidation rate of the Hhy-only mutant was 3-fold higher than the Huc-only strain (**Figure 3b**). H_2_ oxidation was also observed in the cytochrome *bcc-aa*_*3*_ oxidase only strain, showing the complex receives electrons from H_2_ oxidation. However, this activity was 3-fold lower than observed for wild-type cells, suggesting that cytochrome *bd* complex also receives electrons from H_2_ oxidation at 3-days post-OD_max_ (**Figure 3b**). In striking contrast, H_2_ oxidation in the cytochrome *bd* only strain was 6.3-fold greater than wild type (**Figure 3b**). This may be due to an increase in the amount of hydrogenases present in the cells or deregulation of their activity, due to metabolic remodeling to cope with the loss of the proton pumping cytochrome *bcc-aa*_*3*_ oxidase (35). The O_2_ consumption for the wild-type and Huc- and Hhy-only strains, when spiked with H_2_, fit approximately with the 2:1 stoichiometry observed in our initial experiment (**Figure 3c**). However, O_2_ consumption of either oxidase mutants in the presence of H_2_ was significantly higher than wild-type and did not conform to a 2:1 ratio (**Figure 3c**), suggesting H_2_ is co-metabolised with other substrates (e.g. carbon reserves). Taken together, these data show that both terminal oxidase complexes accept electrons from H_2_ oxidation.

Next, we probed the specifics of coupling between Huc and Hhy and the terminal oxidases, by inhibiting the cytochrome *bcc*-*aa*_3_ complex with zinc azide. In wild-type cells, the addition of azide led to a 4.6-fold reduction in H_2_ oxidation, demonstrating that hydrogenase activity is primarily coupled to the cytochrome *bcc-aa*_*3*_ complex at 3-days post-OD_max_. For the Huc-only strain, the addition of zinc azide largely abolished H_2_ oxidation (8.4-fold reduction), suggesting that Huc is obligately coupled to the cytochrome *bcc-aa*_*3*_ complex (**Figure 3b)**. In contrast, only a 1.5-fold reduction in Hhy activity was observed; this demonstrates that while Hhy utilises the cytochrome *bcc*-*aa*_3_ oxidase, it is promiscuous and can also donate electrons to the alternative cytochrome *bd* complex (**Figure 3b**). The addition of azide to the cytochrome *bcc-aa_3_* only strain led near complete inhibition of H_2_ oxidation (10.5-fold decrease) and O_2_ consumption (22.9-fold decrease) (**Figure 3c**). This confirms that Huc and Hhy require an active terminal oxidase to oxidise H_2_, and thus are obligately coupled to the respiratory chain. The high level of H_2_ oxidation observed in the cytochrome *bd* only strain was unchanged by azide treatment, which is expected given this complex is unaffected by azide inhibition (42).

### Huc and Hhy input electrons into the respiratory chain via the quinone pool

Having firmly established that Huc and Hhy activity is coupled to terminal oxidase activity under the conditions tested, we sought to better understand this relationship. To do so, we measured H_2_ oxidation of the wild-type, Huc-only, and Hhy-only strains in the presence of selective respiratory chain inhibitors and uncouplers. First, we tested whether the electrons generated by Hhy and Huc are transferred to the electron carrier menaquinone, which donates electrons to both terminal oxidases in mycobacteria (43). To do so, we tested the effect of HQNO, a competitive inhibitor of quinone-binding (42, 44), on H_2_ oxidation in our wild-type, Huc- and Hhy-only strains. Addition of HQNO led to a 8.6-fold and 10.4-fold decrease in H_2_ oxidation by the Huc- and Hhy-only mutants respectively, demonstrating transport of electrons generated by these enzymes occurs via the menaquinone pool (**Figure 4a,b**). There was a 2.5-fold decrease in the activity of wild-type cells treated with HQNO, confirming that H_2_ oxidation is also menaquinone dependent in a non-mutant background (**Figure 4c**).

Next, we tested the effect of valinomycin on H_2_ oxidation. Valinomycin is an ionophore that binds K^+^ to form a positively charged complex and specifically transporting K^+^ ions across the cellular membrane along the electrical gradient, collapsing the electrical potential component of the proton-motive force (PMF) in respiratory bacteria (45). In *M. smegmatis* at an external of pH above 5, the majority of the PMF is driven by electrical potential (46) and thus addition of valinomycin under our assay conditions leads to a dramatic reduction in the PMF. Huc and Hhy exhibited strikingly different responses to valinomycin. Valinomycin reduced H_2_ oxidation by 20-fold in the Huc-only strain, but increased oxidation by 1.3-fold in the Hhy-only and wild-type strains, compared to untreated cells (**Figure 4**). The near complete loss of Huc activity due to valinomycin treatment indicates this enzyme is energy-dependent, requiring the largely intact PMF to function; this may indicate that the complex is obligately associated with the proton-translocating cytochrome *bcc*-*aa*_3_ supercomplex. Conversely, the increase in Hhy activity demonstrates that this enzyme does not require the PMF, with the increase in H_2_ oxidation possibly resulting from increased metabolic flux as the cells attempt to maintain their membrane potential.

**Figure 4.**
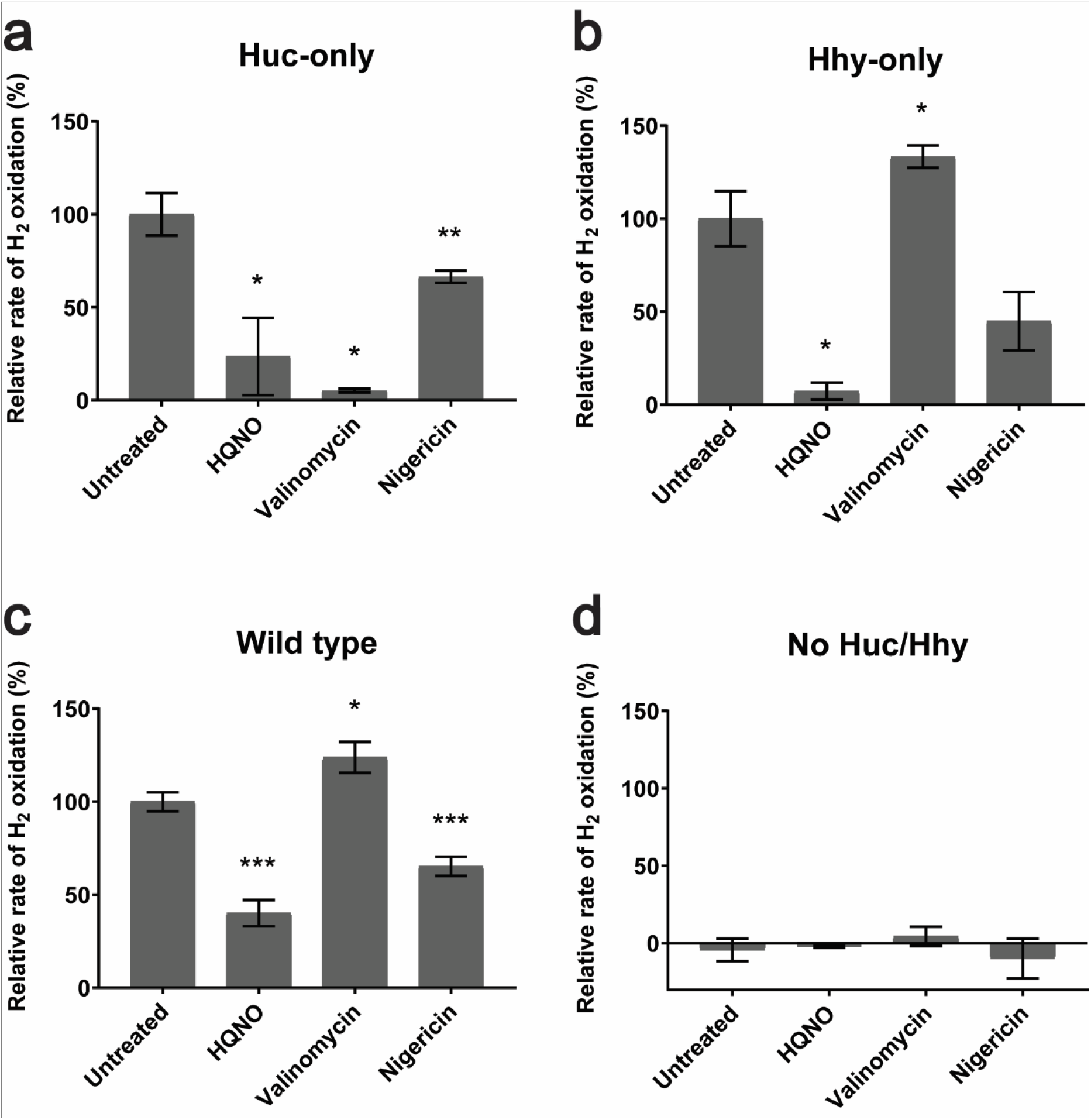
Inhibition of Huc and Hhy coupling to the electron transport chain. H_2_ oxidation rates of (**a**) Huc-only, (**b**) Hhy-only, (**c**) wild-type, and (**d**) no Huc/Hhy (triple hydrogenase deletion) cultures were measured using a hydrogen microelectrode before and after treatment with different respiratory chain uncouplers and inhibitors: HQNO (40 μM), valinomycin (10 μM), nigericin (10 μM). Rates were normalized to mg total protein and expressed as percentage relative to the average rate of untreated cells. Cultures during carbon limitation (3 days post OD_max_ ~3.0) were used. Values with asterisks indicate activity rates that are significantly different from the untreated whole cells based on student’s t-test (* p ≤ 0.05; ** p ≤ 0.01; *** p ≤ 0.001)

Next, we tested the effect of nigericin on H_2_ oxidation. Nigericin is an ionophore which acts as an antiporter of K^+^ and H^+^ ions and is uncharged in its ion bound forms (45). In our assay, nigericin leads to the net efflux of K^+^ ions from and influx of H^+^ ions into the cell, dissipating the proton gradient but not effecting membrane potential. The experimentally determined pH of the external media in our assay was 5.8 (due to media acidification during growth). Under these conditions, ΔpH across the membrane accounts for approximately one third of the PMF in *M. smegmatis* (46). Thus, the addition of nigericin will lead to a significant net influx of protons and acidification of the cytoplasm, shifting the equilibrium for H_2_ towards reduction. The addition of nigericin inhibited H_2_ oxidation to a moderate degree in our assay. Wild-type and Huc-only strains exhibited a ~1.5-fold decrease in activity, while Hhy activity was reduced 2.3-fold (**Figure 4**). The inhibitory effect of nigericin towards Hhy is in contrast with the stimulatory effect of valinomycin on this enzyme. This possibly reflects that, in contrast to valinomycin, nigericin may cause intracellular acidification in addition to diminishing the PMF.

### A model for the integration of hydrogenases in the mycobacterial respiratory chain

Based on the findings from our work we propose a model for integration of Huc and Hhy into the mycobacterial respiratory chain, which we outline in **Figure 5**. Our data show that both Huc and Hhy are obligately coupled to O_2_ reduction via the terminal oxidases of the respiratory chain. Huc preferentially donates electrons to the cytochrome *bcc*-*aa*_3_ complex, while Hhy donates electrons both the cytochrome *bcc*-*aa*_3_ and cytochrome *bd* complexes. Both enzymes are membrane associated, positioning them for the transfer of electrons produced by H_2_ oxidation to the respiratory chain. The size of Huc and its co-migration with the cytochrome *bcc*-*aa*_3_ complex on a native gel suggest that Huc association with the membrane may be mediated by protein-protein interactions, possibly with the terminal oxidase. Inhibition of Huc and Hhy activity by the quinone analogue HQNO suggests that electrons from these enzymes are transferred to the terminal oxidases via the menaquinone pool. It remains to be resolved whether electron transfer to menaquinone is directly mediated by the hydrogenases or through an intermediate protein, for example the FeS proteins co-transcribed with the hydrogenase large and small subunits in the *huc* and *hhy* operons (27). Collapse of the PMF by valinomycin treatment leads to near complete inhibition of Huc activity, but enhancement of Hhy activity. This suggests a distinct relationship exists between these enzymes and the PMF, with Huc requiring an intact PMF, potentially due to its obligate coupling to the cytochrome *bcc*-*aa*_3_ complex.

**Figure 5.**
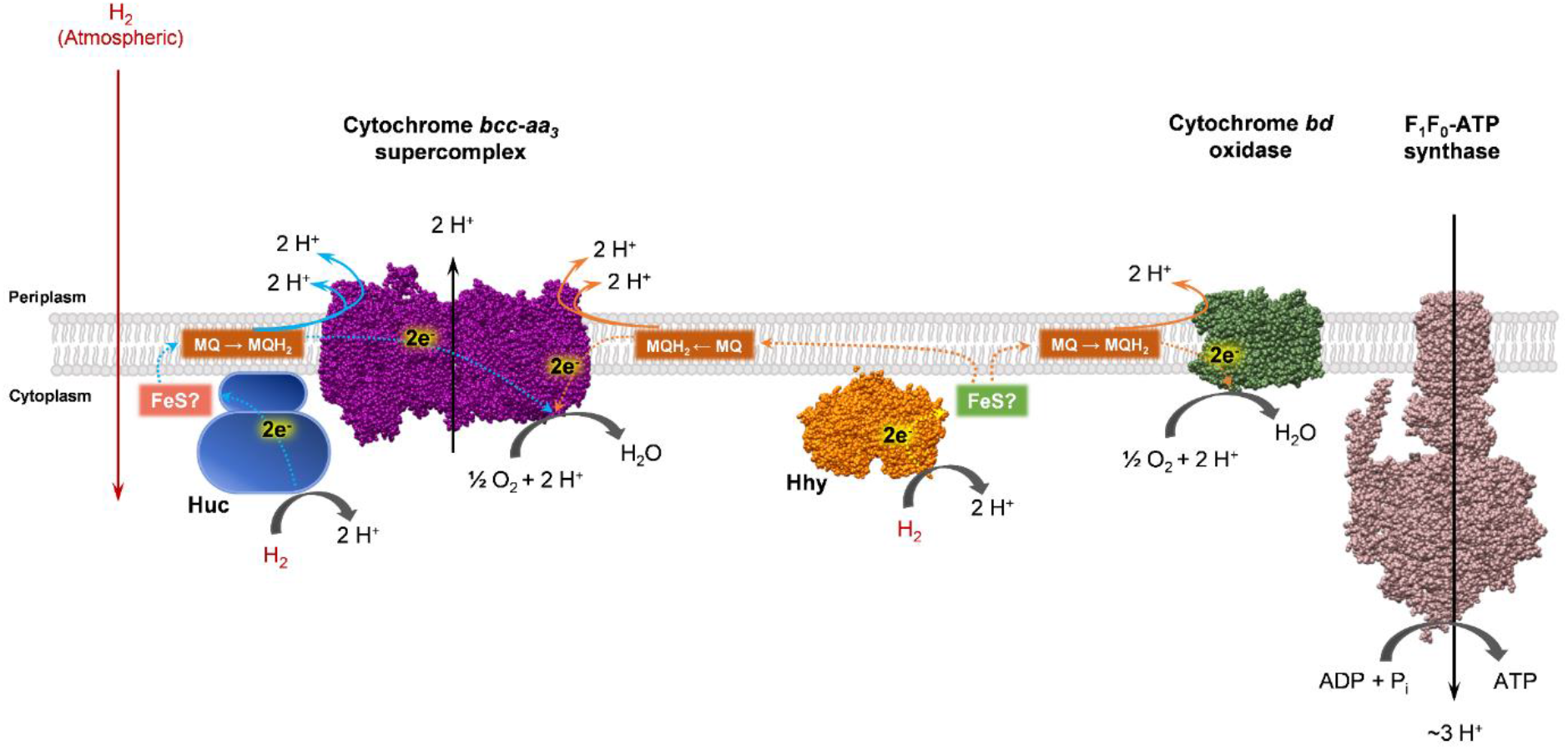
Huc and Hhy differentially energize the mycobacterial respiratory chain during carbon starvation. Both Huc and Hhy oxidise H_2_ to 2 H^+^ and 2e^−^. The electrons are used to reduce membrane-soluble menaquinone (MQ) to menaquinol (MQH_2_). It is possible that the genetically-associated iron-sulfur proteins (FeS) HucE (MSMEG_2268) and HhyE (MSMEG_2718) relay electrons from the hydrogenase to the menaquinone pool. Huc-reduced MQH_2_ transfers electrons exclusively to the cytochrome *bcc*-*aa*_3_ super complex, where they are transferred to the terminal electron acceptor O_2_, yielding H_2_O and resulting in the efflux of 6 H^+^ from the cell. Under starvation or hypoxia, Hhy-derived MQH_2_ transfers electrons to either cytochrome *bcc*-*aa*_3_ or the alternate cytochrome *bd* complex. This results in the efflux of 6 H^+^ or 2 H^+^ from the cell respectively, together with the reduction of O_2_ to H_2_O. The proton gradient generated by H_2_ oxidation maintains membrane potential and allows for the generation of ATP via F_1_F_o_-ATP synthase.

Our data also demonstrates that Huc and Hhy are differentially regulated during mycobacterial growth and persistence. The tightly controlled expression and activity of Huc during the transition between growth and dormancy suggests that it oxidises H_2_ mixotrophically as heterotrophic energy sources become scarce. The proton-motive force generated by Huc, through obligate interaction with the cytochrome *bcc*-*aa*_3_ complex, may help to energise the cell during the transition to dormancy. Expression of *hhyL,* the gene encoding the large Hhy subunit, is upregulated at the cessation of cell division. However, high levels of Hhy-mediated H_2_ oxidation are only observed a number of weeks into dormancy. This suggests that the enzyme primarily functions to meet maintenance needs during persistence, a role which is further supported by its promiscuous utilisation of cytochrome *bd* oxidase. The observed lag between *hhyL* transcription and Hhy activity suggests that regulation of this hydrogenase also occurs downstream of transcription. This possibly provides flexibility to *M. smegmatis,* by limiting synthesis of this resource*-*intensive protein immediately upon exhaustion of carbon-derived energy sources, but allowing rapid deployment of Hhy if resources remain scarce.

While this work provides the basis for understanding how Huc and Hhy are regulated and integrated into cellular metabolism, further investigation is required to fully understand the mechanisms that regulate these enzymes and the biochemistry of their H_2_ oxidation. For example, what are the regulatory pathways that allow the cell to rapidly switch on Huc when resources become scarce and then off again as *M. smegmatis* commits to dormancy? Analogously, how do non-replicating *M. smegmatis* cells regulate transcription and then activity of Hhy during a state of resource limitation? In addition, the data we present suggests physical interactions between Huc and respiratory chain components. Purification of both Huc and Hhy from their native context in the *M. smegmatis* cell will likely provide insight into the protein-protein interactions that mediate electron transfer from these enzymes. Furthermore, purification of these complexes, combined with structural and spectroscopic analysis, will likely provide insight into the mechanisms that underpin the high H_2_ affinity and O_2_ tolerance of Huc and Hhy. In conjunction with this study, these data will provide a richer picture of how mycobacteria consume H_2_ during growth and persistence.

## Materials and Methods

### Bacterial strains and growth conditions

*Mycobacterium smegmatis* mc^2^155 and its derivatives were routinely grown in lysogeny broth (LB) agar plates supplemented with 0.05% (w/v) Tween 80 (LBT) (47). In broth culture, the strain was grown in either LBT or Hartmans de Bont (HdB) minimal medium (pH 7.0) supplemented with 0.2% (w/v) glycerol, 0.05% (w/v) tyloxapol, and 10 mM NiSO_4_. *Escherichia coli* was maintained in LB agar plates and grown in LB broth (48). Selective LB or LBT media used for cloning experiments contained gentamycin at 5 μg mL^−1^ for *M. smegmatis* and 20 μg mL^−1^ for *E. coli*. Cultures were routinely incubated at 37°C, with rotary shaking at 150 rpm for liquid cultures, unless otherwise specified. The strains of *M. smegmatis* and its derivatives and *E. coli* are listed in **Table S2**.

### Insertion of StrepII tags

To facilitate visualization of the hydrogenases in western blots, a StrepII tag was inserted at the C-terminal end of the small subunits of Huc (MSMEG_2262, *hucS*) and Hhy (MSMEG_2720, *hhyS*) through allelic exchange mutagenesis as described previously (33). Two allelic exchange constructs, *hucS*_*StrepII*_ (2656 bp) and *hhyS*_*StrepII*_ (3000 bp), were synthesized by GenScript. These were cloned into the SpeI site of the mycobacterial shuttle plasmid pX33 to yield the constructs pHuc-StrepII and pHhy-StrepII (**Table S3**). The constructs were propagated in *E. coli* TOP10 and transformed into wild-type *M. smegmatis* mc^2^155 cells by electroporation. Gentamycin was used in selective solid and liquid medium to propagate pX33. To allow for permissive temperature-sensitive vector replication, transformants were incubated on LBT gentamycin plates at 28°C until colonies were visible (5-7 days). Resultant catechol-positive colonies were subcultured onto fresh LBT gentamycin plates and incubated at 40°C for 3-5 days to facilitate integration of the recombinant plasmid, via flanking regions, into the chromosome. The second recombination event was facilitated by subculturing catechol-reactive and gentamycin-resistant colonies onto LBT agar plates supplemented with 10% sucrose (w/v) and incubating at 40°C for 3-5 days. Catechol-unreactive colonies were subsequently screened by PCR to distinguish wild-type revertants from Huc-StrepII and Hhy-StrepII mutants. Primers used for screening are listed in **Table S3**.

### Cellular fractionation for detection of Huc and Hhy

The untagged and StrepII-tagged Huc and Hhy were constitutively produced by growing *M. smegmatis* wild-type, Huc-only, Hhy-only, Huc-StrepII, and Hhy-StrepII strains in HdB with 0.2% glycerol as the sole carbon source (24, 47). Cells were grown at 37°C with agitation and harvested by centrifugation (15 min, 10,000 *g*, 4°C) after 1 day post-OD_max_ (~3.0) for wild-type, Huc-only, and Huc-StrepII or 3 days post OD_max_ (~3.0) for wild-type, Hhy-only, and Hhy-StrepII. They were washed in lysis buffer (50 mM Tris-Cl, pH 8.0, 2 mM MgCl_2_, 1 mM PMSF, 5 mg mL^−1^ lysozyme, 40 μg ml^−1^ DNase) and resuspended in the same buffer in a 1:5 cell mass to buffer ratio. The cell suspension was homogenized using a Dounce homogenizer and passed through a cell disruptor (40 Kpsi, four times; Constant Systems One Shot). After removal of unbroken cells by low-speed centrifugation (20 min, 8,000 *g*, 4°C), the whole-cell lysates of wild-type, Huc-only, and Hhy-only strains were used for activity staining. Cell lysates of Huc-StrepII and Hhy-StrepII strains were separated by ultracentrifugation (60 min, 150,000 *g*, 4°C) into cytosol and membrane fractions. Membranes were washed in lysis buffer and analysed by western blotting. Protein concentrations were measured by the bicinchoninic acid method against bovine serum albumin standards.

### Hydrogenase activity staining

Twenty micrograms of each whole-cell lysates was loaded onto two native 7.5% (w/v) Bis-Tris polyacrylamide gels prepared as described elsewhere (49) and run alongside a protein standard (NativeMark Unstained Protein Standard, ThermoFisher Scientific) for 3 h at 25 mA. For total protein determination, gels were stained overnight at 4°C with gentle agitation using AcquaStain Protein Gel Stain (Bulldog Bio). To determine hydrogenase activity, a duplicate gel was incubated in 50 mM potassium phosphate buffer (pH 7.0) supplemented with 500 μM nitroblue tetrazolium chloride (NBT) in an anaerobic jar amended with an anaerobic gas mixture (5% H_2_, 10% CO_2_ 85% N_2_ v/v) overnight at room temperature. Bands present after incubation corresponded to hydrogenase activity.

### Membrane solubilization and western blots

Membrane solubilization was performed by resuspending washed membranes (to a final protein concentration of 1 mg mL^−1^) in solubilization buffer containing 50 mM Tris-Cl pH 8.0, 1 mM PMSF, and 5% (w/v) sodium cholate (50). The solutions were incubated at room temperature with gentle agitation for 3 h. Detergent-soluble proteins were separated from the insoluble material by ultracentrifugation. As a control, membrane was suspended in buffer without sodium cholate. Total proteins in the fractions were visualized in SDS-PAGE and Huc_StrepII and Hhy_StrepII by western blotting. For western blotting, 20 μg total protein was loaded and ran on to Bolt™ 4-12% SDS polyacrylamide gels after boiling in Bolt™ SDS sample buffer and 50 mM dithiothreitol. The proteins in the gels were then transferred onto PVDF membrane using Trans-Blot® SD Semi-Dry Transfer Cell (Bio-Rad) set at 15 V for 60 m. Following transfer, the protein-containing PVDF membrane was blocked with 3% (w/v) bovine serum albumin in phosphate-buffered saline (PBS), pH 7.4 with 0.1% (v/v) Tween 20 (PBST). PVDF membrane was washed three times in 20 mL of PBST and finally resuspended in 10 mL of the same buffer. Strep-Tactin horse radish peroxidase (HRP) conjugate was then added at a 1:100,000 dilution. Peroxide-mediated chemilluminescence of luminol catalyzed by the HRP was developed according to manufacturer’s specifications (Amersham ECL Prime detection reagent, GE Life Sciences) and the Strep-Tactin(HRP conjugated)–StrepII-tag complex was visualized in a Fusion Solo S (Fischer Biotech) chemiluminescence detector.

### Gene expression analysis

For qRT-PCR analysis, five synchronized sets of biological triplicate cultures (30 mL) of wild-type *M. smegmatis* were grown in 125 mL aerated conical flasks. Each set of triplicates was quenched either at OD_600_ 0.3, OD_600_ 1.0, 1-day post-OD_max_ (OD_600_ ~3.0), 3-days post-OD_max_, or 3-weeks post-OD_max_ with 60 mL cold 3:2 glycerol:saline solution (−20°C). They were subsequently harvested by centrifugation (20,000 × *g*, 30 minutes, −9°C), resuspended in 1 mL cold 1:1 glycerol:saline solution (−20°C), and further centrifuged (20,000 × *g*, 30 minutes, −9°C). For cell lysis, pellets were resuspended in 1 mL TRIzol Reagent, mixed with 0.1 mm zircon beads, and subjected to five cycles of bead-beating (4,000 rpm, 30 seconds) in a Biospec Mini-Beadbeater. Total RNA was subsequently extracted by phenol-chloroform extraction as per manufacturer’s instructions (TRIzol Reagent User Guide, Thermo Fisher Scientific) and resuspended in diethylpyrocarbonate (DEPC)-treated water. RNA was treated with DNase using the TURBO DNA-free kit (Thermo Fisher Scientific) as per the manufacturer’s instructions. cDNA was then synthesized using SuperScript III First-Strand Synthesis System for qRT-PCR (Thermo Fisher Scientific) with random hexamer primers as per the manufacturer’s instructions. qPCR was used to quantify the levels of the target genes *hucL* (Huc) and *hhyL* (Hhy) and housekeeping gene *sigA* against amplicon standards of known concentration. All reactions were run in a single 96-well plate using the PowerUp SYBR Green Master Mix (Thermo Fisher Scientific) and LightCycler 480 Instrument (Roche) according to each manufacturers’ instructions. A standard curve was created based on the cycle threshold (Ct) values of *hucL*, *hhyL*, and *sigA* amplicons that were serially diluted from 10^8^ to 10 copies (R^2^ > 0.98). The copy number of the genes in each sample was interpolated based on each standard curve and values were normalized to *sigA* expression. For each biological replicate, all samples, standards, and negative controls were run in technical duplicate. Primers used in this work are summarized in **Table S3**.

### Microrespiration measurements

Rates of H_2_ oxidation or O_2_ consumption were measured amperometrically according to previously established protocols (27, 30). For each set of measurements, either a Unisense H_2_ microsensor or Unisense O_2_ microsensor electrode were polarised at +800 mV or −800 mV, respectively, with a Unisense multimeter. The microsensors were calibrated with either H_2_ or O_2_ standards of known concentration. Gas-saturated phosphate-buffered saline (PBS; 137 mM NaCl, 2.7 mM KCl, 10 mM Na_2_HPO_4_ and 2 mM KH_2_PO_4_, pH 7.4) was prepared by bubbling the solution with 100% (v/v) of either H_2_ or O_2_ for 5 min. In uncoupler/inhibitor-untreated cells, H_2_ oxidation was measured in 1.1 mL microrespiration assay chambers. These were amended with 0.9 mL cell suspensions of *M. smegmatis* wild-type or derivative strains either at OD_600_ 0.3, OD_600_ 1.0, 1-day post-OD_max_ (OD_600_ ~3.0), 3-days post-OD_max_, or 3-weeks post-OD_max_, They were subsequently amended with 0.1 mL H_2_-saturated PBS and 0.1 mL O_2_-saturated PBS. Chambers were stirred at 250 rpm, room temperature. For cells at mid-stationary phase, following measurements of untreated cells, the assay mixtures were treated with either 10 μM nigericin, 10 μM valinomycin, 40 μM *N*-oxo-2-heptyl-4-hydroxyquinoline (HQNO), or 250 μM zinc azide before measurement. In O_2_ consumption measurements, initial O_2_ consumption without the addition of H_2_ were measured in microrespiration assay chambers sequentially amended with 0.9 mL cell suspensions of *M. smegmatis* wild-type or derivative strains at mid-stationary phase (3-days post-OD_max_) and 0.1 mL O_2_-saturated PBS (0.1 mL) with stirring at 250 rpm, room temperature. After initial measurements, 0.1 mL of H_2_-saturated PBS was added into the assay mixture and changes in O_2_ concentrations were recorded. Additionally, O_2_ consumption was measured in cytochrome *bcc-aa*_*3*_-only strains treated with 250 μM zinc azide. In both H_2_ and O_2_ measurements, changes in concentrations were logged using Unisense Logger Software (Unisense, Denmark). Upon observing a linear change in either H_2_ or O_2_ concentration, rates of consumption were calculated over a period of 20 s and normalised against total protein concentration.

## Acknowledgements

This work was supported by an ARC DECRA Fellowship (DE170100310; awarded to C.G.), an NHMRC New Investigator Grant (APP5191146; awarded to C.G.), a Monash University Science-Medicine Seed Grant (awarded to C.G. and M.J.C.), and a Monash University Doctoral Scholarship (awarded to P.R.F.C.). We thank Dr Ralf Schittenhelm and Dr Cheng Huang of the Monash Proteomic & Metabolic Facility for performing mass spectrometry analyses.

## Footnotes

### Author contributions

C.G. and G.M.C. conceived the study. C.G., P.R.F.C., G.M.C., R.G., K.H., M.J.C., and C.G.W. designed experiments. P.R.F.C. performed experiments. P.R.F.C., C.G., R.G., K.H., and G.M.C. analysed data. P.R.F.C., R.G., and C.G. wrote the paper with input from all authors.

### Competing financial interests

The authors declare no competing financial interests.

## References

1. Rhee, T. S., Brenninkmeijer, C. A. M., and Rockmann, T. (2006) The overwhelming role of soils in the global atmospheric hydrogen cycle. Atmos. Chem. Phys. 6, 1611–1625

2. Schmidt, U. (1974) Molecular hydrogen in the atmosphere. Tellus A. 26, 78–90

3. Constant, P., Chowdhury, S. P., Pratscher, J., and Conrad, R. (2010) Streptomycetes contributing to atmospheric molecular hydrogen soil uptake are widespread and encode a putative high-affinity [NiFe]-hydrogenase. Environ. Microbiol. 12, 821–829

4. Greening, C., Carere, C. R., Rushton-Green, R., Harold, L. K., Hards, K., Taylor, M. C., Morales, S. E., Stott, M. B., and Cook, G. M. (2015) Persistence of the dominant soil phylum *Acidobacteria* by trace gas scavenging. Proc. Natl. Acad. Sci. 112, 10497–10502

5. Islam, Z. F., Cordero, P. R. F., Feng, J., Sean, Y. C., Thanavit, K. B., Gleadow, R. M., Carere, C. R., Stott, M. B., Chiri, E., and Greening, C. (2019) Two Chloro fl exi classes independently evolved the ability to persist on atmospheric hydrogen and carbon monoxide. ISME J. 13, 1801–1813

6. Greening, C., Villas-Bôas, S. G., Robson, J. R., Berney, M., and Cook, G. M. (2014) The growth and survival of *Mycobacterium smegmatis* is enhanced by co-metabolism of atmospheric H_2_. PLoS One. 9, e103034

7. Liot, Q., and Constant, P. (2016) Breathing air to save energy - new insights into the ecophysiological role of high-affinity [NiFe]-hydrogenase in Streptomyces avermitilis. Microbiologyopen. 5, 47–59

8. Meredith, L. K., Rao, D., Bosak, T., Klepac-Ceraj, V., Tada, K. R., Hansel, C. M., Ono, S., and Prinn, R. G. (2014) Consumption of atmospheric hydrogen during the life cycle of soil-dwelling actinobacteria. Environ. Microbiol. Rep. 6, 226–238

9. Constant, P., Poissant, L., and Villemur, R. (2009) Tropospheric H_2_ budget and the response of its soil uptake under the changing environment. Sci. Total Environ. 407, 1809–1823

10. Ehhalt, D. H., and Rohrer, F. (2009) The tropospheric cycle of H_2_: A critical review. Tellus, Ser. B Chem. Phys. Meteorol. 61, 500–535

11. Piché-Choquette, S., Khdhiri, M., and Constant, P. (2018) Dose-response relationships between environmentally-relevant H_2_ concentrations and the biological sinks of H_2_, CH_4_ and CO in soil. Soil Biol. Biochem. 123, 190–199

12. Bay, S., Ferrari, B., and Greening, C. (2018) Life without water: How do bacteria generate biomass in desert ecosystems? Microbiol. Aust. 39, 28–32

13. Greening, C., Biswas, A., Carere, C. R., Jackson, C. J., Taylor, M. C., Stott, M. B., Cook, G. M., and Morales, S. E. (2016) Genomic and metagenomic surveys of hydrogenase distribution indicate H_2_ is a widely utilised energy source for microbial growth and survival. ISME J. 10, 761–777

14. Ji, M., Greening, C., Vanwonterghem, I., Carere, C., Bay, S., Steen, J., Montgomery, K., Lines, T., Beardall, J., Dorst, J. van, Snape, I., Stott, M., Hugenholtz, P., and Ferrari, B. (2017) Atmospheric trace gases support primary production in Antarctic desert ecosystems. Nature. 552, 400–403

15. Kanno, M., Constant, P., Tamaki, H., and Kamagata, Y. (2015) Detection and isolation of plant-associated bacteria scavenging atmospheric molecular hydrogen. Environ. Microbiol. 18, 2495–2506

16. Kessler, A. J., Chen, Y.-J., Waite, D. W., Hutchinson, T., Koh, S., Popa, M. E., Beardall, J., Hugenholtz, P., Cook, P. L. M., and Greening, C. (2019) Bacterial fermentation and respiration processes are uncoupled in anoxic permeable sediments. Nat. Microbiol. 4, 1014–1023

17. Khdhiri, M., Hesse, L., Popa, M. E., Quiza, L., Lalonde, I., Meredith, L. K., Röckmann, T., and Constant, P. (2015) Soil carbon content and relative abundance of high affinity H_2_-oxidizing bacteria predict atmospheric H_2_ soil uptake activity better than soil microbial community composition. Soil Biol. Biochem. 85, 1–9

18. Lynch, R. C., Darcy, J. L., Kane, N. C., Nemergut, D. R., and Schmidt, S. K. (2014) Metagenomic evidence for metabolism of trace atmospheric gases by high-elevation desert actinobacteria. Front. Microbiol. 5, 1–13

19. Piché-Choquette, S., Khdhiri, M., and Constant, P. (2017) Survey of high-affinity H_2_-oxidizing bacteria in soil reveals their vast diversity yet underrepresentation in genomic databases. Microb. Ecol. 74, 771–775

20. Schwartz, E., Fritsch, J., and Friedrich, B. (2013) H_2_-Metabolizing Prokaryotes. in The Prokaryotes: Prokaryotic Physiology and Biochemistry (Rosenberg, E., DeLong, E. F., Lory, S., Stackebrandt, E., and Thompson, F. eds), pp. 119–199, Springer, Berlin/Heidelberg, Germany

21. Ferrari, B. C., Bissett, A., Snape, I., Dorst, J. Van, Palmer, A. S., Ji, M., Siciliano, S. D., Stark, J. S., Winsley, T., and Brown, M. V (2016) Geological connectivity drives microbial community structure and connectivity in polar, terrestrial ecosystems. 18, 1834–1849

22. Price, P. B., and Sowers, T. (2004) Temperature dependence of metabolic rates for microbial growth, maintenance, and survival. Proc. Natl. Acad. Sci. 101, 4631–4636

23. Greening, C., Constant, P., Hards, K., Morales, S. E., Oakeshott, J. G., Russell, R. J., Taylor, M. C., Berney, M., Conrad, R., and Cookb, G. M. (2015) Atmospheric hydrogen scavenging: From enzymes to ecosystems. Appl. Environ. Microbiol. 81, 1190–1199

24. Greening, C., Berney, M., Hards, K., Cook, G. M., and Conrad, R. (2014) A soil actinobacterium scavenges atmospheric H_2_ using two membrane-associated, oxygen-dependent [NiFe] hydrogenases. Proc. Natl. Acad. Sci. 111, 4257–4261

25. Schäfer, C., Friedrich, B., and Lenza, O. (2013) Novel, oxygen-insensitive group 5 [NiFe]-hydrogenase in Ralstonia eutropha. Appl. Environ. Microbiol. 79, 5137–5145

26. Schäfer, C., Bommer, M., Hennig, S. E., Jeoung, J. H., Dobbek, H., and Lenz, O. (2016) Structure of an Actinobacterial-Type [NiFe]-Hydrogenase Reveals Insight into O_2_-Tolerant H_2_ Oxidation. Structure. 24, 285–292

27. Berney, M., Greening, C., Hards, K., Collins, D., and Cook, G. M. (2014) Three different [NiFe] hydrogenases confer metabolic flexibility in the obligate aerobe Mycobacterium smegmatis. Environ. Microbiol. 16, 318–330

28. Constant, P., Poissant, L., and Villemur, R. (2008) Isolation of *Streptomyces* sp. PCB7, the first microorganism demonstrating high-affinity uptake of tropospheric H_2_. ISME J. 2, 1066–1076

29. Myers, M. R., and King, G. M. (2016) Isolation and characterization of *Acidobacterium ailaaui* sp. nov., a novel member of Acidobacteria subdivision 1, from a geothermally heated Hawaiian microbial mat. Int. J. Syst. Evol. Microbiol. 66, 5328–5335

30. Carere, C. R., Hards, K., Houghton, K. M., Power, J. F., Mcdonald, B., Collet, C., Gapes, D. J., Sparling, R., Boyd, E. S., Cook, G. M., Greening, C., and Stott, M. B. (2017) Mixotrophy drives niche expansion of verrucomicrobial methanotrophs. ISME J. 11, 2599–2610

31. Berney, M., Greening, C., Conrad, R., Jacobs, W. R., and Cook, G. M. (2014) An obligately aerobic soil bacterium activates fermentative hydrogen production to survive reductive stress during hypoxia. Proc. Natl. Acad. Sci. 111, 11479–11484

32. Berney, M., and Cook, G. M. (2010) Unique flexibility in energy metabolism allows mycobacteria to combat starvation and hypoxia. PLoS One. 5, e8614

33. Tran, S. L., and Cook, G. M. (2005) The F1F0-ATP Synthase of Mycobacterium smegmatis Is Essential for Growth. J. Bacteriol. 187, 5023–5028

34. Cook, G. M., Hards, K., Vilchèze, C., Hartman, T., and Berney, M. (2013) Energetics of Respiration and Oxidative Phosphorylation in Mycobacteria. Microbiol. Spectr. 2, 1–30

35. Wiseman, B., Nitharwal, R. G., Fedotovskaya, O., Schäfer, J., Guo, H., Kuang, Q., Benlekbir, S., Sjöstrand, D., Ädelroth, P., Rubinstein, J. L., Brzezinski, P., and Högbom, M. (2018) Structure of a functional obligate complex III2IV2 respiratory supercomplex from Mycobacterium smegmatis. Nat. Struct. Mol. Biol. 25, 1128–1136

36. Borisova, V. B., Gennis, R. B., Hemp, J., and Verkhovsky, M. I. (2011) The cytochrome bd respiratory oxygen reductases. Biochim. Biophys. Acta. 1807, 1398–1413

37. Safarian, S., Rajendran, C., Müller, H., Preu, J., Langer, J. D., Hirose, T., Kusumoto, T., Sakamoto, J., Michel, H., and Ovchinnikov, S. (2016) Structure of a bd oxidase indicates similar mechanisms for membrane-integrated oxygen reductases. Science (80-.). 352, 583–586

38. Cordero, P. R. F., Bayly, K., Leung, P. M., Huang, C., Islam, Z. F., Schittenhelm, R. B., King, G. M., and Greening, C. (2019) Atmospheric carbon monoxide oxidation is a widespread mechanism supporting microbial survival. ISME J. https://doi.org/10.1038/s41396-019-0479-8

39. Piché-Choquette, S., and Constanta, P. (2019) Molecular hydrogen, a neglected key driver of soil biogeochemical processes. Appl. Environ. Microbiol. 85, 1–19

40. Kim, M., Jang, J., Binte, N., Rahman, A. B., Pethe, X. K., and Berry, X. E. A. (2015) Isolation and Characterization of a Hybrid Respiratory Supercomplex Consisting of Mycobacterium tuberculosis Cytochrome bcc and Mycobacterium smegmatis Cytochrome aa3. J. Biol. Chem. 290, 14350–14360

41. Lu, P., Heineke, M. H., Koul, A., Andries, K., Cook, G. M., Lill, H., Van Spanning, R., and Bald, D. (2015) The cytochrome bd-type quinol oxidase is important for survival of Mycobacterium smegmatis under peroxide and antibiotic-induced stress. Sci. Rep. 5, 10333:1–10

42. Cook, G. M., Greening, C., Hards, K., and Berney, M. (2014) Energetics of Pathogenic Bacteria and Opportunities for Drug Development. in Advances in Microbial Physiology (Poole, R. K. ed), pp. 1–65, Academic Press, Cambridge, Massachusetts, USA

43. Dhiman, R. K., Mahapatra, S., Slayden, R. A., Boyne, M. E., Lenaerts, A., Hinshaw, J. C., Angala, S. K., Chatterjee, D., Biswas, K., Narayanasamy, P., Kurosu, M., and Crick, D. C. (2009) Menaquinone synthesis is critical for maintaining mycobacterial viability during exponential growth and recovery from non-replicating persistence. Mol. Microbiol. 72, 85–97

44. Matsuno-Yagi, A., and Hatefi, Y. (2002) Ubiquinol-Cytochrome c Oxidoreductase. J. Biol. Chem. 271, 6164–6171

45. Nicholls, D. G., and Ferguson, S. J. (2013) Ion transport across energy-conserving membranes. in Bioenergetics, 4th Ed., pp. 13–25, Academic Press, Cambridge, Massachusetts, USA

46. Rao, M., Streur, T. L., Aldwell, F. E., and Cook, G. M. (2001) Intracellular pH regulation by Mycobacterium smegmatis and Mycobacterium bovis BCG. Microbiology. 147, 1017–1024

47. Gebhard, S., Tran, S. L., and Cook, G. M. (2006) The Phn system of *Mycobacterium smegmatis*: a second high-affinity ABC-transporter for phosphate. Microbiology. 152, 3453–3465

48. Green, M. R., and Sambrook, J. (2012) Molecular cloning: a laboratory manual, 4th Ed., Cold Spring Harbor Laboratory Press, Cold Spring Harbor, New York, USA

49. Walker, J. M. (2009) Nondenaturing Polyacrylamide Gel Electrophoresis of Proteins. in The Protein Protocols Handbook, 3rd Ed., pp. 171–176, Humana Press, Totowa, New Jersey, USA

50. Heikal, A., Nakatani, Y., Dunn, E., Weimar, M. R., Day, C. L., Baker, E. N., Lott, J. S., Sazanov, L. A., and Cook, G. M. (2014) Structure of the bacterial type II NADH dehydrogenase: A monotopic membrane protein with an essential role in energy generation. Mol. Microbiol. 91, 950–964

51. Snapper, S. B., Melton, R. E., Mustafa, S., Kieser, T., and Jr, W. R. J. (1990) Isolation and characterization of efficient plasmid transformation mutants of Mycobacterium smegmatis. Mol. Microbiol. 4, 1911–1919

52. Lu, X., Williams, Z., Hards, K., Tang, J., Cheung, C. Y., Aung, H. L., Wang, B., Liu, Z., Hu, X., Lenaerts, A., Woolhiser, L., Hastings, C., Zhang, X., Wang, Z., Rhee, K., Ding, K., Zhang, T., and Cook, G. M. (2019) Pyrazolo[1,5- a]pyridine Inhibitor of the Respiratory Cytochrome bcc Complex for the Treatment of Drug-Resistant Tuberculosis. ACS Infect. Dis. 5, 239–249

53. Gebhard, S., Tran, S. L., and Cook, G. M. (2006) The Phn system of Mycobacterium smegmatis: A second high-affinity ABC-transporter for phosphate. Microbiology. 152, 3453–3465

