## Supplementary Material for "Two uptake hydrogenases differentially interact with the aerobic respiratory chain during mycobacterial growth and persistence"

### Supporting information

**Figure S1. Hydrogenase activity staining in blue native PAGE.** Differential native activity staining (left panel) of Huc and Hhy in whole-cell lysates of *M. smegmatis* WT strain harvested at either 1-day post-$OD_{max}$ (for Huc staining) or 3-days post-$OD_{max}$ (for Hhy staining). The high molecular weight species (>700 kDa) in the Huc activity stain (pointed by an arrow) is consistent with the band seen in Figure 2a. The corresponding coomassie stain of the total protein is shown at the right panel.

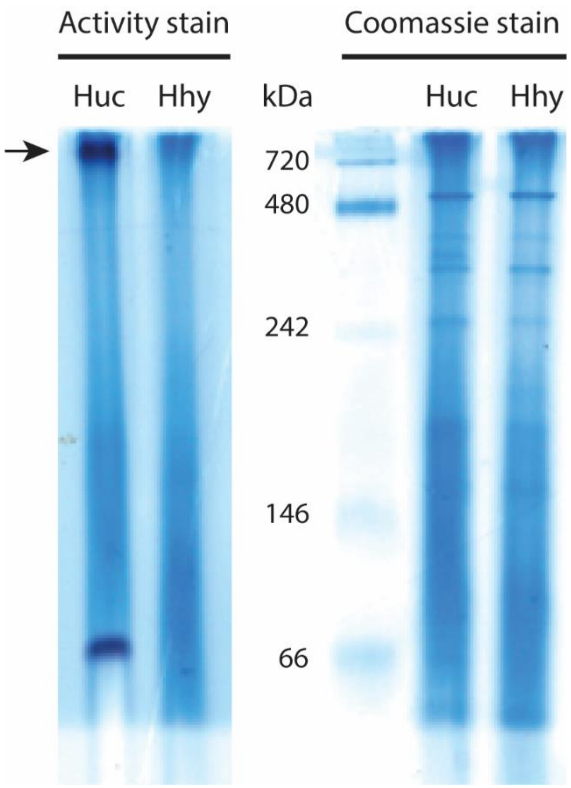

**Table S1. Selected proteins identified in Huc and Hhy activity stain bands through mass spectrometry.**

| Protein ID | Protein | Log Probability | Coverage (%) |
|---|---|---|---|
| Proteins in the "Huc" band | | | |
| YP_886615.1 | HucL | 27.27 | 15.89 |
| YP_888539.1 | QcrA | 234.92 | 58.09 |
| YP_888540.1 | QcrB | 168.88 | 32.97 |
| YP_888538.1 | QcrC | 41.91 | 22.01 |
| YP_888712.1 | CtaA | 7.75 | 4.12 |
| YP_888545.1 | CtaB | 193.51 | 45.75 |
| YP_888537.1 | CtaC | 13.50 | 6.90 |
| Proteins in the "Hhy" band | | | |
| YP_887054.1 | HhyS | 25.86 | 8.83 |
| YP_887053.1 | HhyL | 206.78 | 39.46 |

**Table S2. Bacterial strains and plasmids used in this study.**

| Strain or plasmid | Description | Source or reference |
|---|---|---|
| **Strains** | | |
| $mc^2155$ | *Mycobacterium smegmatis* wild type strain | (51) |
| Huc-StrepII | $mc^2155$ with StrepII tag inserted at the C-terminus of MSMEG_2262 (*hucS-$_{StrepII}$-hucL)* | This study |
| Hhy-StrepII | $mc^2155$ with StrepII tag inserted at the C-terminus of MSMEG_2720 (*hhyS-$_{StrepII}$-hhyL)* | This study |
| Huc-only | $mc^2155$ with markerless deletions of MSMEG_2719, MSMEG_3931 (Δ*hhyL*Δ*hyhS*) | (27) |
| Hhy-only | $mc^2155$ with markerless deletions of MSMEG_2262, MSMEG_3931 Δ*hucS*Δ*hyhS*) | (27) |
| No Huc/Hhy | $mc^2155$ with markerless deletions of MSMEG_2262, MSMEG_2719, MSMEG_3931 (Δ*HucS*Δ*HhyL*Δ*hyhS*) | (27) |
| cyt *bcc-aa$_3$* -only | $mc^2155$ with markerless deletions of cytochrome *bd* subunits (Δ*cydAB*) | (52) |
| cyt *bd*-only | $mc^2155$ with markerless deletions of cytochrome *bcc-aa$_3$* subunits (Δ*qcrCAB*) | Gregory M. Cook (Otago University, NZ) |
| TOP10 | *Escherichia coli* strain F- *mcrA* Δ( *mrr-hsd*RMS-*mcr*BC) Φ80*lac*ZΔM15 Δ *lac*X74 *rec*A1 *ara*D139 Δ(*araleu*)7697 *gal*U *gal*K *rps*L (StrR) *end*A1 *nup*G | ThermoFischer |
| **Plasmids** | | |
| pX33 | $Gm^r$, *sacB*, mycobacterial Ts *ori*, p15A *ori*, *xylE* | (53) |
| pHuc_StrepII | 2656 bp *hucS$_{-StrepII}$-hucL* fragment in pX33 | This study |
| pHhy_StrepII | 3000 bp *hhyS$_{-StrepII}$-hhyL* fragment in pX33 | This study |

**Table S3. List of primers used in this work.**

| Purpose | Primer name | Sequence |
|---|---|---|
| Construction of pHuc_StrepII | HucF (SpeI restriction site) | AAAACTAGTATGGCATCGGTGCTTTGGTTC |
| | HucR (SpeI restriction site) | AAAACTAGTTCACACCATCCCGTTGATCAC |
| Construction of pHhy_StrepII | HhyF (SpeI restriction site) | AAAACTAGTATGCCAACGGAGGCTGCAGT |
| | HhyR (SpeI restriction site) | AAAACTAGTTCAGTCCCCGGTCGCGGA |
| Sequencing of constructs to verify cloning of insert | T7F | TAATACGACTCACTATAGGG |
| | pX33_SpeIR | AATAGATCATCGTCGCCG |
| | HucInt1 | TACATGGGTCTGGCGGCGG |
| | HucInt2 | CCAGGGCGTGGGCAACTAC |
| | HhyInt1 | CACGGTGGCGGCATCGCG |
| | HhyInt2 | GCTGGGGTGCTGGGGTTC |
| Screening of Huc-StrepII and Hhy-StrepII insertion mutants | Huc_chromF | AATCCGGCAGCAGCCCTG |
| | Huc_chromR | GACCGACCCCGTCGTCAC |
| | Hhy_chromF | GCGTCTTCACTCGGGACG |
| | Hhy_chromR | GTCAACGGGAACGTCGGC |
| | StrepR | GAACTGCGGGTGCGACCA |
| qRT-PCR of *hucL* (MSMEG_2263) | q2263F | ACCCTGATCCGCAACATCTG |
| | q2263R | GTCGTTCCAATGCTCCAGGA |
| qRT-PCR of *hhyL* (MSMEG_2719) | q2719F | AATCAGCACACCAACCCCAA |
| | q2719F | CCTCGAACTTCTCCCACGTC |
| qRT-PCR of *sigA* (MSMEG_2758) | qsigAF | CTCAACGCCGAAGAAGAGGT |
| | qsigAR | GCCCTTGGTGTAGTCGAACT |
